# PhosF3C: A Feature Fusion Architecture with Fine-Tuned Protein Language Model and Conformer for prediction of general phosphorylation site

**DOI:** 10.1101/2024.12.25.630296

**Authors:** Yuhuan Liu, Haitian Zhong, Jixiu Zhai, Xueying Wang, Tianchi Lu

**Affiliations:** Cuiying Honors College, Lanzhou University, 222 South Tianshui Road, Lanzhou 730000, China; New Laboratory of Pattern Recognition (NLPR), State Key Laboratory of Multimodal Artificial Intelligence Systems (MAIS), Institute of Automation, Chinese Academy of Sciences (CASIA); School of Mathematics and Statistics, Lanzhou University, 222 South Tianshui Road, Lanzhou 730000, China; Department of Computer Science, City University of Hong Kong, Kowloon, Hong Kong; Department of Computer Science, City University of Hong Kong (Dongguan), Dongguan 523000, China

**Keywords:** Protein Phosphorylation, Large language model, LoRA, Conformer

## Abstract

Protein phosphorylation, a key post-translational modification (PTM), provides essential insight into protein properties, making its prediction highly significant. Using the emerging capabilities of large language models (LLMs), we apply LoRA fine-tuning to ESM2, a powerful protein large language model, to efficiently extract features with minimal computational resources, optimizing task-specific text alignment. Additionally, we integrate the conformer architecture with the Feature Coupling Unit (FCU) to enhance local and global feature exchange, further improving prediction accuracy. Our model achieves state-of-the-art (SOTA) performance, obtaining AUC scores of 79.5%, 76.3%, and 71.4% at the S, T, and Y sites of the general data sets. Based on the powerful feature extraction capabilities of LLMs, we conduct a series of analyses on protein representations, including studies on their structure, sequence, and various chemical properties (such as Hydrophobicity (GRAVY), Surface Charge, and Isoelectric Point). We propose a test method called Linear Regression Tomography (LRT) which is a top-down method using representation to explore the model’s feature extraction capabilities, offering a pathway to improved interpretability.

## 1: Introduction

Protein phosphorylation is one of the most crucial post-translational modifications (PTMs). It plays an essential role in regulating various cellular processes. These processes include signal transduction, cell cycle control, metabolism, apoptosis, and protein-protein interactions. Phosphorylation occurs through the addition of a phosphate group to specific amino acids, typically serine, threonine, or tyrosine. This modification can cause structural changes in proteins. It can also modulate enzymatic activity or affect binding affinity and localization within the cell. Phosphorylation is particularly significant in cellular signaling pathways. It acts as a molecular on/off switch to control key processes such as cell division, immune responses, and gene expression (1, 2).

Phosphorylation’s role is especially prominent in cancer. Aberrant activation of kinase-driven pathways, such as EGFR and HER2, leads to uncontrolled cell proliferation and tumor growth. This makes kinase inhibitors a vital class of targeted therapies (3, 4). In neurodegenerative diseases, abnormal phosphorylation of proteins like tau contributes to pathological aggregates. These findings offer insights into therapeutic approaches (5, 6). In cardiovascular diseases, phosphorylation of regulatory proteins affects heart muscle contractility. Targeting these pathways holds potential for treating heart failure (7, 8). Given its extensive involvement in disease mechanisms, predicting phosphorylation sites is crucial. This helps identify biomarkers, understand pathological processes, and design effective therapeutic interventions, especially for complex diseases like cancer, neurodegenerative disorders, and heart disease (9).

In recent years, the study of PTMs, particularly phosphorylation, has advanced significantly. This progress is due to the development of high-throughput technologies and machine learning models. These advancements have enabled the identification and mapping of phosphorylation sites at an unprecedented scale. New approaches, such as mass spectrometry and phosphoproteomics, are pivotal in understanding the dynamic nature of phosphorylation. These methods reveal its variations across different cell types and conditions (10–13). Computational methods have also evolved. Recent models achieve greater accuracy in predicting phosphorylation sites and their roles in various diseases. These developments provide deeper insights into how phosphorylation regulates cellular functions and contributes to disease mechanisms (14, 15).

Several models have demonstrated notable performance in protein phosphorylation prediction. Each model has unique strengths and limitations. **DeepPSP** performs well by utilizing both global and local information for complete protein sequences. However, its effectiveness decreases when analyzing protein fragments, as it struggles without full sequence context (16). **MusiteDeep** uses rotational attention to extract dual-dimensional sequence features. Despite this, its reliance on one-hot encoding results in feature loss during embedding, limiting its ability to capture complex sequence information (17). **PtransIPS** excels in feature extraction with its dual-embedding approach. It combines sequence and structural information using ProtTrans (18) and EMBER2 (19). However, it faces significant challenges in computation time and resource requirements when processing large datasets (20). The rapid evolution of LLMs has provided powerful tools for protein phosphorylation site prediction. These models leverage emergent capabilities to automatically extract complex features from protein sequences. They capture subtle relationships between amino acids and phosphorylation sites (21). LLMs deliver enhanced performance in feature extraction and deep contextual understanding, making them more efficient and accurate than traditional methods. Fine-tuning techniques, such as LoRA, further improve their efficiency. This positions LLMs as a promising approach for advancing phosphorylation prediction (22, 23).

Based on the protein language model ESM2 (24), our model addresses the limitations of existing approaches by achieving highly efficient feature extraction. It improves the prediction of short protein sequence segments while significantly reducing storage and computational resource requirements. This is accomplished through LoRA fine-tuning, ensuring task alignment with minimal computational overhead. The model is further enhanced with the conformer architecture (25), which captures both global and local features. FCU is used to facilitate dynamic interactions between these features, an aspect often overlooked by previous models. Additionally, we applied our model to other protein-related tasks, further demonstrating the high generalizability of this framework. Across various tasks, the model showed strong feature extraction capabilities, proving its effectiveness and adaptability in multiple protein prediction challenges.

Lastly, we conducted a series of studies on protein representations generated by ESM2, applying traditional machine learning methods to analyze specific features. Using techniques such as random forests (26) and multiple linear regression (27), we developed a top-down analysis approach called LRT. Inspired by RePE (28, 29), this method identifies the principal directions of properties within the representations. This approach offers a potential pathway for improving model interpretability.

## 2: Materials and Methods

### A. Dataset

#### A.1 Dataset overview

Our dataset is divided into two parts: **general datasets** and **task-specific datasets**.

##### General Datasets

The general dataset used in our study is a comprehensive fusion of data sourced from MusiteDeep and DeepPSP. The dataset integrate phosphorylation site information from key databases including UniPort/SWISSPROT(30), PhosphoSitePlus(31), (32), and Phospho.ELM(33). This dataset includes general phosphorylation, including kinase-specific data, providing a rich resource for phosphorylation prediction tasks.

In terms of structure, although we incorporate kinase-specific information from several families (CDK, MAPK, CK2, PKA, PKC, AGC, CMGC, CAMK), we opted not to explicitly train separate models for each kinase family, in contrast to methods used in some other studies. This approach helps to maintain model generalization across different types of phosphorylation events while still benefiting from the rich diversity provided by kinase-specific information.

The dataset was then split into training and testing sets. For evaluation, a portion of the dataset was also reserved for independent testing. These methods offer a robust way to measure model performance across various conditions.

In addition to the general datasets, we utilized two specific datasets for phosphorylation site prediction in special tasks: **DeepIPS**(34) and **PhosAF**(35).

##### DeepIPS Dataset

The DeepIPS dataset consists of phosphorylation sites from human A549 cells infected with SARS-CoV-2, comprising data from the literature. Protein sequences were truncated into 33-residue segments centered on serine (S), threonine (T), or tyrosine (Y). Positive samples were defined as phosphorylated, while non-phosphorylated segments were treated as negative samples.

##### PhosAF Dataset

The PhoSAF dataset contains phosphorylation sites from human proteins, sourced from UniProt/Swiss-Prot, PhosphoSitePlus, and Phospho.ELM. CD-HIT(36) was used to ensure that the similarity of sequence was less than 40%. Protein fragments centered on serine (S), threonine (T), or tyrosine (Y) were extracted.

The detailed information and sample proportions for each dataset, including the number of positive and negative samples across Serine (S), Threonine (T), and Tyrosine (Y) phosphorylation sites, are presented in Table 1. The overall dataset composition and the positive and negative sample ratios for the merged general dataset, PhosAF, and DeepIps datasets are visually summarized in Figure 1.

**Table 1.**
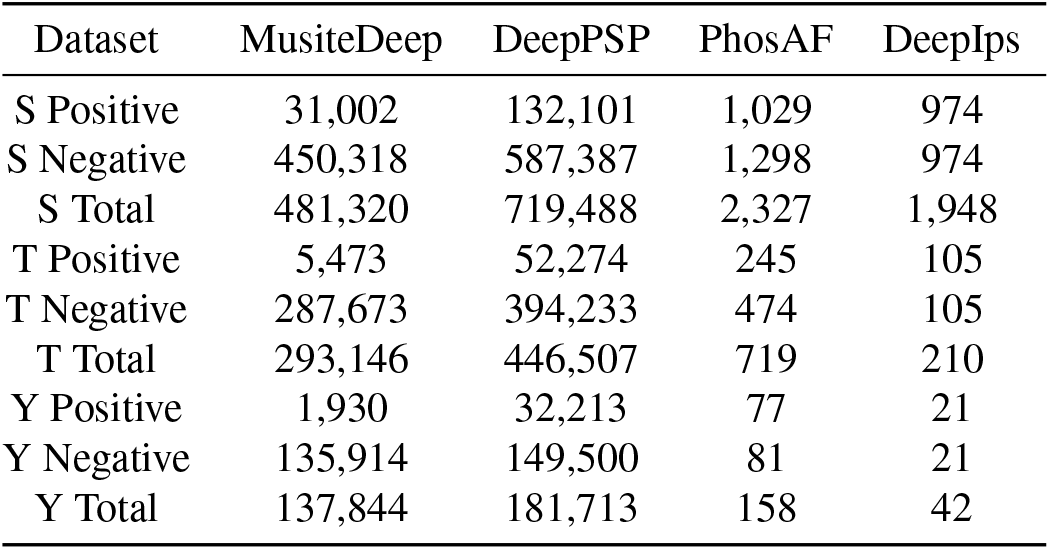
Overview of phosphorylation site data across different datasets.

**Table 2.** Training data overview for Serine (S), Threonine (T), and Tyrosine (Y) phosphorylation sites.

| Train Set | Serine (S) | Threonine (T) | Tyrosine (Y) |
| --- | --- | --- | --- |
| Positive | 122074 | 47122 | 29303 |
| Negative | 580626 | 395141 | 167830 |
| Total | 702700 | 442263 | 197133 |

**Fig 1.**
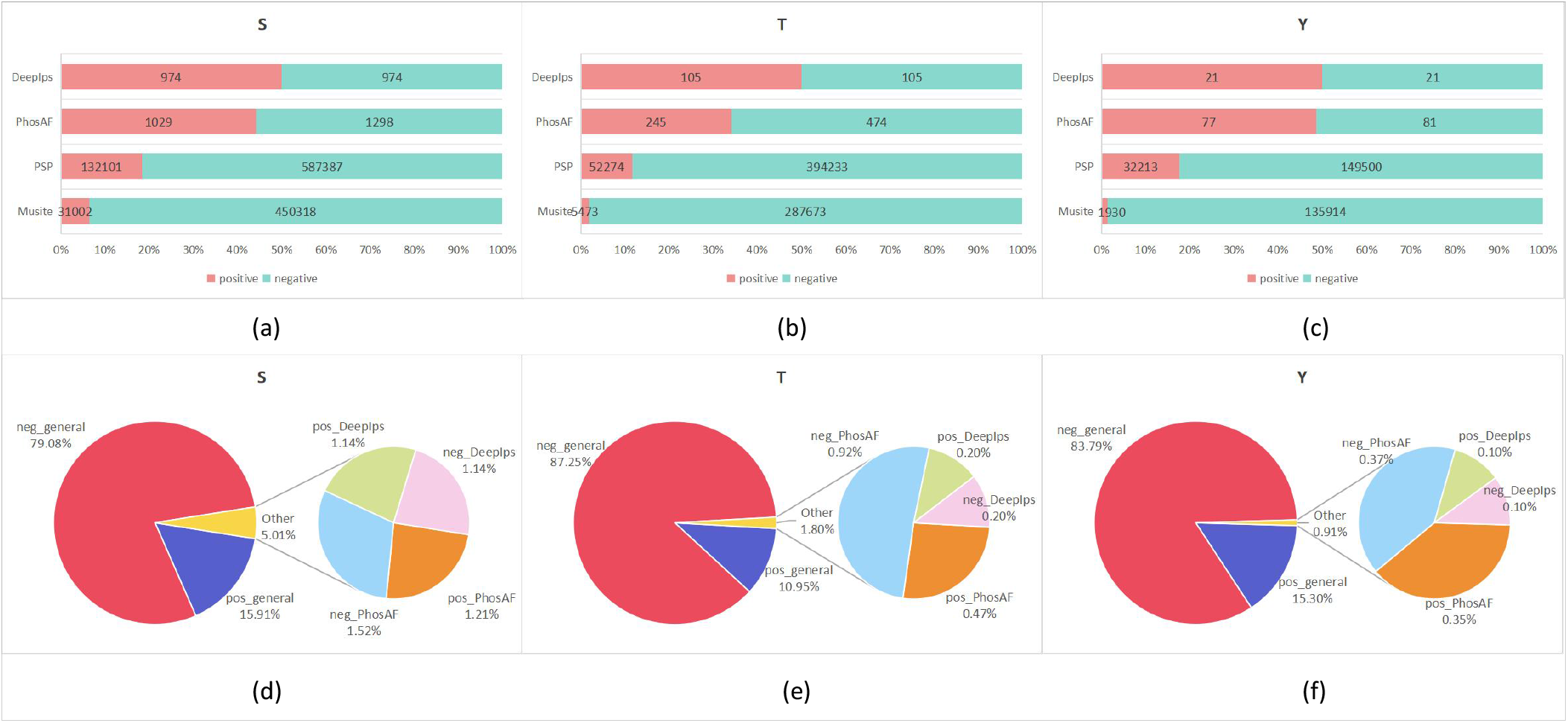
Dataset overview showing the number of Serine (S), Threonine (T), and Tyrosine (Y) phosphorylation sites and positive-to-negative ratios in individual datasets (a), (b), (c). (d), (e), (f) display the distribution and proportion of samples in the combined test set and task-specific datasets, including positive and negative ratios.

**Fig 2.**
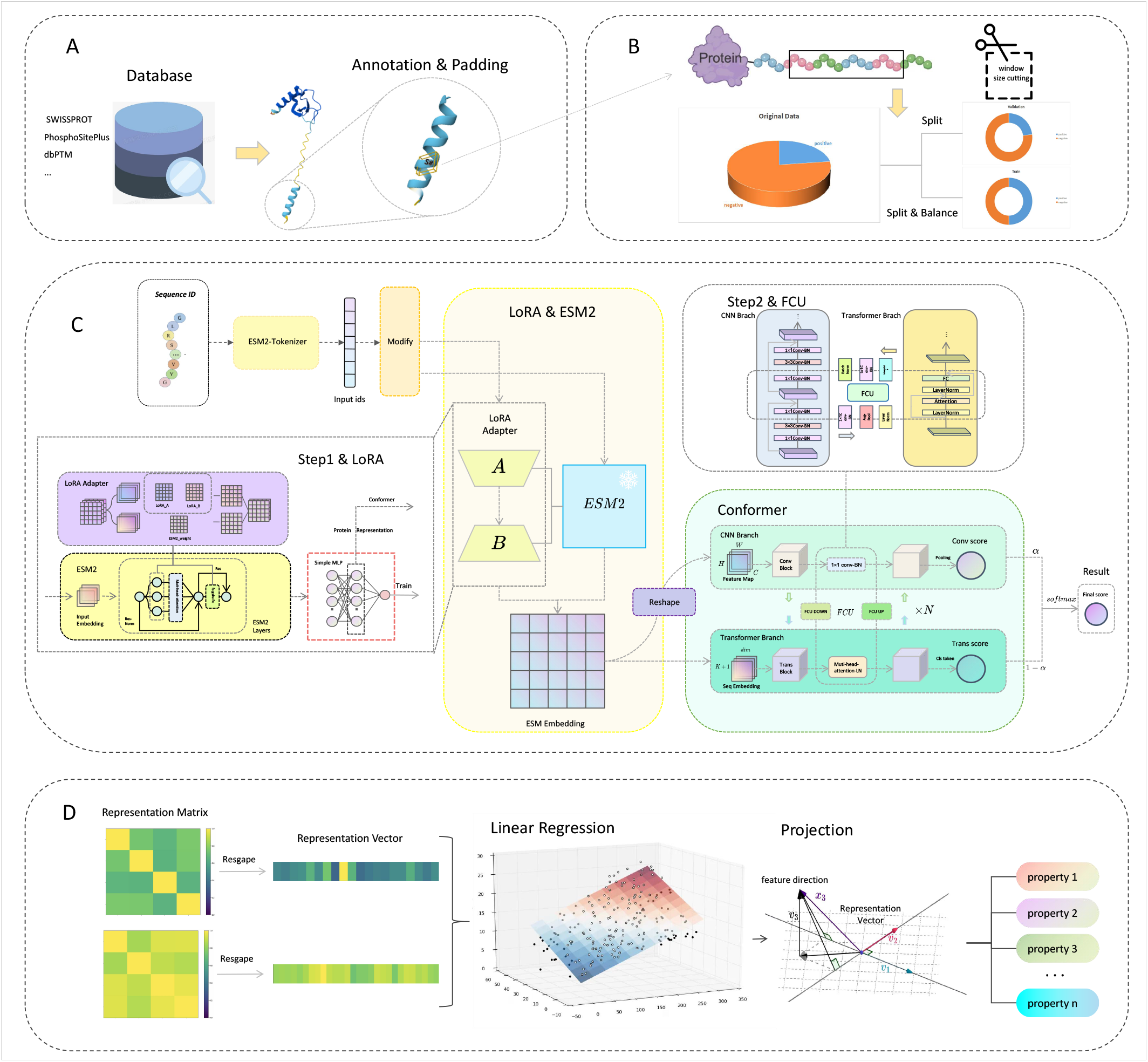
A and B show the data extraction, preprocessing, and generation of training and validation sets. B specifically handles the extraction of window-based phosphorylation site counts from labeled proteins, creating the training and validation sets with balanced data for the training set. C illustrates the model architecture and training process: Step 1 involves fine-tuning ESM2 with LoRA for task alignment, and Step 2 combines the fine-tuned ESM2 with the downstream Conformer model, where two branches perform separate predictions and FCU facilitates information interaction between them. D demonstrates the LRT process, where the feature matrix is first transformed into feature vectors, linear regression is applied to identify the principal directions of properties, and the projection shows how well the model extracts these properties.

**Fig 3.**
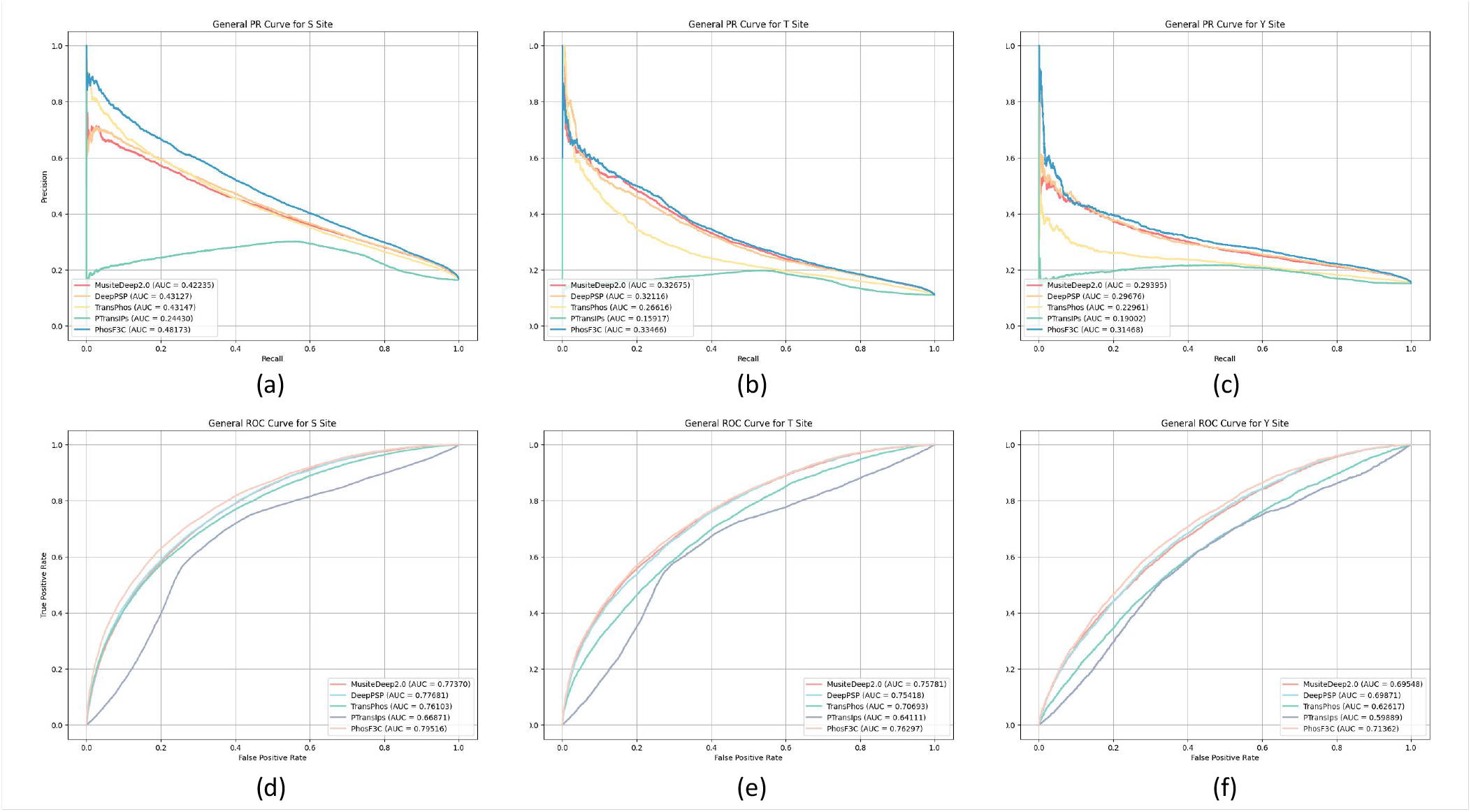
Figures (a), (b), and (c) present the PR curves for Serine (S), Threonine (T), and Tyrosine (Y) sites on the general test dataset, comparing the performance of the PhosF3C model with other currently well-performing protein phosphorylation prediction models. Figures (d), (e), and (f) show the corresponding ROC curves for the same sites.

#### A.2 Dataset Splitting Training and Testing

After merging the general datasets, we randomly split the dataset into a general training set and a general test set. Specifically, we isolated 1500 independent(10% of the whole general dataset), full-length sequences to form the general test set, while the remaining sequences were used as the general training set. For the other two datasets, PhosAF and DeepIPS, we used only the test portions that were originally defined in their respective datasets.

Regarding the training set, due to the class imbalance where negative samples far outnumber positive samples, we applied a strategy to balance the dataset by repeating the entire positive sample set until the number of positive samples exceeds the number of negative samples, and then truncating the positive samples to match the number of negative samples. This ensures an equal number of positive and negative samples for training.

For model training, we only used the general training set, and the train portions of the original PhosAF and DeepIPS datasets were excluded. This strategy allows us to evaluate the model’s generalization ability across different test sets, emphasizing the robustness and transferability(different task) of the model developed on the general dataset.

Training set information is presented in 2, and test set information is presented in Table 3 and portion in Figure 1.

**Table 3.**
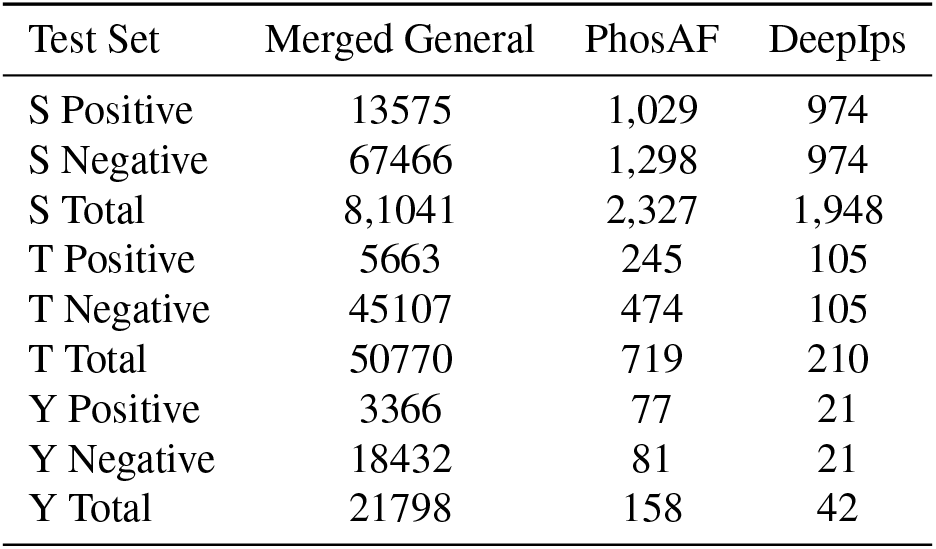
Test data overview for General, PhosAF, and DeepIps datasets.

| Test Set | Merged General | PhosAF | DeepIps |
| --- | --- | --- | --- |
| S Positive | 13575 | 1,029 | 974 |
| S Negative | 67466 | 1,298 | 974 |
| S Total | 8,1041 | 2,327 | 1,948 |
| T Positive | 5663 | 245 | 105 |
| T Negative | 45107 | 474 | 105 |
| T Total | 50770 | 719 | 210 |
| Y Positive | 3366 | 77 | 21 |
| Y Negative | 18432 | 81 | 21 |
| Y Total | 21798 | 158 | 42 |

### B. Method

#### B.1. Method overview

Our approach consists of two main steps designed to enhance protein phosphorylation site prediction through effective feature extraction and integration. Step 1: Initial Fine-Tuning with LoRA We begin by employing LoRA to fine-tune a simple architecture consisting of ESM2 combined with two layers of MLP (multi-layer perceptron). This architecture is specifically used for the protein phosphorylation prediction task. The trained checkpoint is aligned with the identification task, leveraging the self-regressive encoding capabilities of the ESM2 model to capture the abstract features necessary for this task. This enables a rational allocation of weights to the relevant features.

Step 2: Feature Extraction and Integration Next, we reintegrate the obtained checkpoint back into the ESM2 model, using it as a feature extractor for downstream tasks. We connect the Conformer architecture to this setup. The resulting feature matrix is divided into two parts and subjected to appropriate transformations before being processed through the Conformer’s CNN(37) branch and Transformer(38) branch for local and global feature extraction, respectively. Each branch comprises multiple layers of structures. To facilitate effective communication between the branches, we employ FCU for feature fusion, allowing timely information exchange between the two branches. Finally, we derive distinct evaluation parameters from each branch and utilize MLP to obtain predictive assessments for the phosphorylation sites.

#### B.2. LoRA Fine-tuning

In conventional deep learning, the weights obtained through gradient descent enable the model’s predictions to more closely align with the target labels, thereby reducing the loss function. For illustration, let’s consider a multi-layer perceptron (MLP) and denote the forward propagation as model and input as x.

The update rule for the weights can be expressed as:

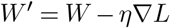

where *W* ^*′*^ is the updated weight, *W* is the original weight, *η* is the learning rate, and *∇ L* is the gradient of the loss function.

After the update, we can expand the updated weight matrix into the original weight part and the gradient descent update part. Let the rank of the gradient be denoted as *r*. Thus, we have:

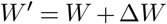

Where

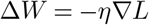

Next, substituting this into the forward propagation gives us:

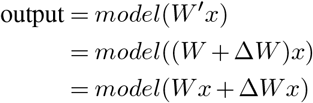

To analyze the change in predictions, we can derive the difference in predicted values before and after the update:

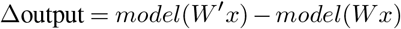

Using the first-order Taylor expansion around *W*, we find:

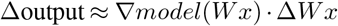

However, due to the substantial number of parameters in LLMs, storing and computing these updates can be resource-intensive. Therefore, we aim for a simpler representation of Δ*W*, proposing that it can be expressed as a product of two lower-rank matrices, *A* and *B*, with ranks *r*_*a*_ and *r*_*b*_, respectively, where both are much smaller than *r*:

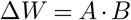

Substituting this back into the forward propagation reveals that these can effectively be represented as two subsequent MLP forward propagations:

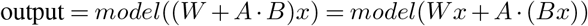

We can formally view *ABx* as *x* passing through two layers of MLP:

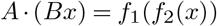

where *f*_1_ and *f*_2_ represent the two layers of the MLP. We denote these two layers of MLP as the **LoRA adapter**.

Based on this concept, the LoRA adapter can be applied in any context where MLPs are utilized, such as in the feedforward networks of transformers, or in the attention mechanisms’ *Q, K, V* matrices, as well as in the final output projection layer.

#### B.3. Conformer

##### CNN Branch

The CNN branch follows a feature pyramid structure where the resolution of feature maps decreases with depth while the number of channels increases. The branch is divided into four stages, each consisting of multiple convolution blocks. Each convolution block contains several bottlenecks. A bottleneck includes a 1×1 down-projection convolution, a 3×3 spatial convolution, a 1×1 up-projection convolution, and a residual connection between input and output. Unlike transformers, which project sequence representation patches into vectors in a single step, potentially losing local details, CNNs apply sliding convolution kernels over feature maps, preserving fine local features. This allows the CNN branch to provide detailed local information for the Transformer branch.

##### Transformer Branch

The Transformer branch, inspired by ViT(39), consists of multiple transformer blocks. Each block includes a multi-head self-attention module and an MLP block, both of which are preceded by LayerNorm and include residual connections. A class token is added to the patch embeddings for classification purposes. Since the CNN branch encodes both local features and spatial information, positional embeddings are not required, allowing the model to handle high-resolution representation more efficiently.

##### FCU

To bridge the gap between the local features from the CNN branch and the global representations from the Transformer branch, we introduce FCU. FCU aligns the dimensionality of the CNN feature maps (C × H × W) with the Transformer patch embeddings ((K + 1) × E, 1 for class token and K for sequence length), using 1×1 convolutions and down-sampling/up-sampling modules (average pooling and interpolating)to ensure compatibility between the branches. FCU progressively closes the semantic gap between the CNN’s local convolutional features and the Transformer’s global self-attention mechanisms, enabling effective feature fusion across all layers except the first.

### C. LRT

#### C.1. Feature Importance via Random Forests

We start by identifying key features that are important for the phosphorylation site prediction task. Using a Random Forest model, we determine the most influential properties of proteins, such as biochemical characteristics like surface charge or hydrophobicity. Once these important features are identified, we obtain their real values (measured experimentally) or predicted values (derived from a specialized prediction model) for a set of proteins.

#### C.2. Representation and Feature Direction

For each protein, we extract its representation using the ESM2. Our objective is to find a principal direction within the representation matrix that best correlates with the specific biochemical property of interest. To do this, we apply **multivariate linear regression** to map the feature values to the corresponding direction in the representation space. Mathematically, this can be formulated as:

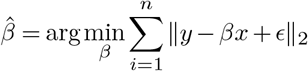

where *y* represents the observed or predicted feature values, *X* is the matrix of ESM2 representations, *β* is the vector representing the direction in the representation space that corresponds to the feature, and *E* is the error term. In our experiments, we use the 128 samples with the highest and lowest feature values to ensure a clear separation of extremes.

#### C.3. Projection and Feature Approximation

After finding the optimal direction *β* through regression, we project the protein’s ESM2 representation onto this direction. The value of the feature for each protein is then approximated as the projection of its representation onto the direction *β*:

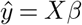

This projected value *ŷ* serves as an estimate of the protein’s biochemical property.

#### C.4. Comparison with Real Distribution

Finally, we compare the distribution of the predicted values *ŷ* with the real or experimentally measured distribution of the feature values. If the feature extraction model (ESM2) is efficient, the distribution of the projected values will closely match the real distribution. This comparison provides a way to measure the model’s ability to extract meaningful features.

### D. Performance Evaluation

In our evaluation, we utilize a range of metrics to assess model performance across different aspects of prediction quality:

- **AUC (Area Under the Curve)**:

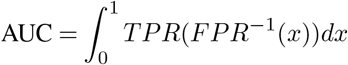
- **Accuracy (ACC)**:

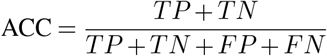
- **Matthews Correlation Coefficient (MCC)**:

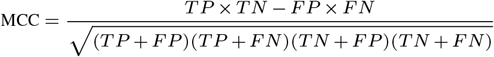

- **F1 Score**:

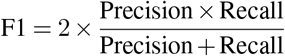
- **Recall (REC)**:

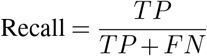
- **Precision (PRE)**:

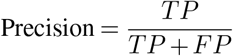

**Definitions:**

- *TP* : True Positive
- *TN* : True Negative
- *FP* : False Positive
- *FN* : False Negative

## 3: Result

### A. Independent test of PhosF3C for phosphorylation site identification

To comprehensively evaluate the performance of our models, we compared them with five established phosphorylation site prediction tools, including DeepPSP, MusiteDeep, TransPhos(40), PtransIPs and PhosF3C. Each model was trained on the general dataset and tested on three independent test datasets, following a consistent training and testing split as described in the *Dataset Splitting Training and Testing* section. It’s important to note that all models were trained with same window size 31 (sequence’s global information was masked during training) and same training pattern. Predictions were evaluated using a phosphorylation determination threshold set at 0.5, ensuring uniformity across comparisons. This threshold selection avoids bias related to sample imbalance and allows for fair model performance assessment across different datasets. Performance on General Dataset The results are presented in Table 4 and 3. For Serine (S) and Threonine (T) sites, the PhosF3C model achieved the best performance across most metrics, with an AUC of 0.7952 for S and 0.763 for T sites. This surpasses other models like DeepPSP and MusiteDeep by margins of 1.84% and 2.15% for S sites, and 0.88% and 0.52% for T sites, respectively. For Tyrosine (Y) sites, although DeepPSP showed competitive performance, the PhosF3C still demonstrated superior AUC (0.7136). Performance on PhosAF Dataset For the PhosAF dataset, PhosF3C achieves the highest overall AUC values for serine (S) and threonine (T) sites, with scores of 0.9155 and 0.8934, respectively. These results are notably higher than the second-best performing models, indicating its superior ability to capture the underlying patterns of these phosphorylation sites. Additionally, it excels in other critical metrics, such as F1 (0.813 for S and 0.7292 for T) and MCC (0.6542 for S and 0.5733 for T), reflecting its balance between precision and recall, as well as its capacity to make accurate predictions. Even for the more challenging tyrosine (Y) site predictions, PhosF3C performs competitively with an AUC of 0.7178 and an MCC of 0.2872.

**Table 4.** Performance index on general dataset categorized by phosphorylation site type.

| Type | Model | AUC | ACC | MCC | F1 | RECALL | PRECISION |
| --- | --- | --- | --- | --- | --- | --- | --- |
| S | DeepPSP | 0.7768 | 0.6703 | 0.3031 | 0.4305 | 0.7441 | 0.3029 |
|  | MusiteDeep2.0 | 0.7737 | 0.6275 | 0.2904 | 0.4167 | 0.7941 | 0.2824 |
|  | TransPhos | 0.761 | 0.6841 | 0.2926 | 0.426 | 0.7 | 0.3062 |
|  | PTransIPS | 0.6687 | 0.6678 | 0.246 | 0.3918 | 0.6519 | 0.28 |
|  | PhosF3C | 0.7952 | 0.7281 | 0.3449 | 0.4648 | 0.705 | 0.3467 |
| T | DeepPSP | 0.7542 | 0.7355 | 0.2453 | 0.336 | 0.5999 | 0.2333 |
|  | MusiteDeep2.0 | 0.7578 | 0.6625 | 0.2387 | 0.3202 | 0.7127 | 0.2065 |
|  | TransPhos | 0.7069 | 0.7537 | 0.1985 | 0.3041 | 0.4824 | 0.222 |
|  | PTransIPS | 0.6411 | 0.6559 | 0.1782 | 0.2822 | 0.6133 | 0.1832 |
|  | PhosF3C | 0.763 | 0.7254 | 0.2591 | 0.3435 | 0.6438 | 0.2342 |
| Y | DeepPSP | 0.6987 | 0.6493 | 0.2099 | 0.3571 | 0.6307 | 0.2491 |
|  | MusiteDeep2.0 | 0.6955 | 0.6515 | 0.2024 | 0.3524 | 0.6141 | 0.2471 |
|  | TransPhos | 0.6262 | 0.6274 | 0.1429 | 0.3137 | 0.5514 | 0.2192 |
|  | PTransIPS | 0.5989 | 0.6018 | 0.135 | 0.3058 | 0.5805 | 0.2076 |
|  | PhosF3C | 0.7136 | 0.6334 | 0.2263 | 0.3661 | 0.6857 | 0.2498 |

Performance on DeepIPs Dataset On the DeepIPs dataset, PhosF3C maintains its strong performance. For serine (S) and threonine (T) sites, it achieves top AUC scores of 0.875 and 0.8584, respectively, which are complemented by solid F1 scores (0.7843 for S and 0.7897 for T). These results indicate that the model not only identifies true positive phosphorylation sites effectively but also minimizes false positives. While the AUC for tyrosine (Y) sites is slightly lower at 0.7302, it remains competitive compared to other models. More detailed test results are provided in Supplementary. This supplementary includes all relevant metrics and further analysis to ensure a comprehensive evaluation of their performance alongside the primary models.

### B. Training with LoRA and FCU improves the performance of PhosF3C

In this section, we present the ablation study to evaluate the effectiveness of key components in our models, specifically focusing on the impact of LoRA and FCU. All experiments were conducted on the general test set.

Our results (Table 5) highlight that the LoRA-enhanced model improves performance by selectively enhancing task-relevant abstract features, thus increasing their contribution to the prediction task. The PhosF3C with LoRA achieves an AUC of 0.7876 for S sites, 0.7606 for T sites, and 0.7011 for Y sites, surpassing the base conformer model (AUC: S: 0.7726, T: 0.7381, Y: 0.6901). The MCC values for LoRA (S: 0.3409, T: 0.2401, Y: 0.2121) are also higher than those of the conformer model (S: 0.2996, T: 0.2016, Y: 0.1976), indicating a stronger correlation with the true labels.

**Table 5.** Performance index on general dataset categorized by phosphorylation site type.

| Type | Model | AUC | ACC | MCC | F1 | RECALL | PRECISION |
| --- | --- | --- | --- | --- | --- | --- | --- |
| S | conformer | 0.7726 | 0.6576 | 0.2996 | 0.4265 | 0.7602 | 0.2964 |
|  | LoRA | 0.7876 | 0.7435 | 0.3409 | 0.4641 | 0.6628 | 0.357 |
|  | PhosF3C | 0.7952 | 0.7281 | 0.3449 | 0.4648 | 0.705 | 0.3467 |
| T | conformer | 0.7381 | 0.5584 | 0.2016 | 0.2855 | 0.7909 | 0.1742 |
|  | LoRA | 0.7606 | 0.6755 | 0.2401 | 0.3232 | 0.6947 | 0.2106 |
|  | PhosF3C | 0.763 | 0.7254 | 0.2591 | 0.3435 | 0.6438 | 0.2342 |
| Y | conformer | 0.6901 | 0.5812 | 0.1976 | 0.3458 | 0.7169 | 0.2279 |
|  | LoRA | 0.7011 | 0.6096 | 0.2121 | 0.356 | 0.6988 | 0.2389 |
|  | PhosF3C | 0.7136 | 0.6334 | 0.2263 | 0.3661 | 0.6857 | 0.2498 |

FCU significantly enhances performance beyond LoRA’s improvements alone. Specifically, PhosF3C with LoRA achieves higher AUCs (S: 0.7952, T: 0.763, Y: 0.7136) compared to LoRA alone (AUC: S: 0.7876, T: 0.7606, Y: 0.7011). The MCC values also reflect this improvement, with the PhosF3C with FCU and LoRA attaining MCC scores of 0.3449 for S sites, 0.2591 for T sites, and 0.2263 for Y sites, compared to LoRA-only MCCs (S: 0.3409, T: 0.2401, Y: 0.2121).

The PR and ROC curves are shown in Figure 4, clearly demonstrating that the combined strengths of LoRA and FCU enhance model performance. Additionally, Figure 4 indicates that LoRA effectively aligns ESM2 feature extraction with the task requirements.

**Fig 4.**
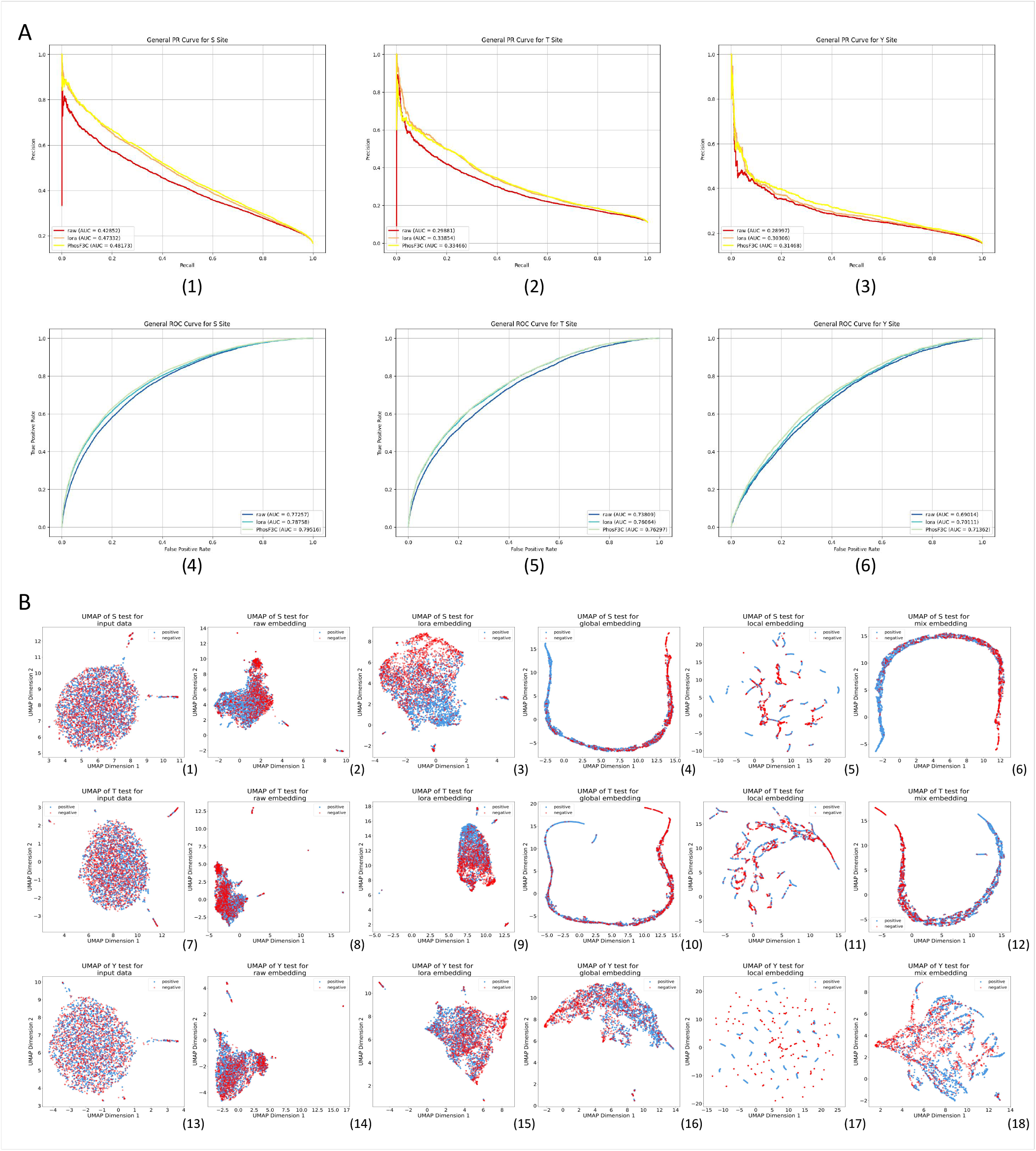
Ablation study for phosphorylation site prediction, showing PR and ROC curves for different model variations: ESM2 (raw checkpoint) + Conformer, ESM2 (finetuned by LoRA) + 2 MLP layers, and ESM2 (finetuned by LoRA) + Conformer. A shows the PR and ROC curves for Serine (S), Threonine (T), and Tyrosine (Y) sites, respectively;B shows the UMAP visualization in a two-dimensional representation during the training process, reflecting the distribution of protein features at different stages in low-dimensional space, with negative and positive samples annotated to highlight classification performance. The stages include: raw (pre-fine-tuned ESM2 embedding), lora (post-LoRA fine-tuning), global (features processed by the transformer branch), local (features processed by the CNN branch), and mix (a combination of both feature processes)..

### C. Visualizing the feature extraction process of PhosF3C with 2D UMAP

To investigate the model’s capability in distinguishing phosphorylation sites we performed a visualization of features extracted during different stages of training using uniform manifold approximation and projection (UMAP)(41)(as show in 4). Initially, embeddings from the ESM model showed limited capacity to differentiate between positive and negative phosphorylation sites, as the samples were largely intermingled and lacked discernible boundaries . This observation indicates that the raw ESM embeddings alone were not capable of filtering out the most relevant abstract features for this task.

However, after incorporating LoRA, we observe a clear improvement in feature alignment, with task-relevant features gradually emerging . This demonstrates that LoRA plays a critical role in filtering these abstract features, leading to a better focus on key elements essential for phosphorylation prediction.

As we progress further, conformer’s FCU exhibits its strength in facilitating the interaction between local and global information. This dual extraction from the CNN and Transformer branches results in a distinct separation of positive and negative samples, with both branches contributing complementary perspectives to improve classification accuracy.

### D. Applying LRT in feature extracting analysis

In the general test, we conducted an experiment using Linear LRT to analyze the ability of our model to capture meaningful physicochemical properties of proteins. First, we predicted various physicochemical properties of the proteins using Biopython(42) over 65 window sizes. For each protein sequence, the values of these properties were distributed, and their importance for phosphorylation site discrimination was evaluated using a random forest classifier. We also calculate the property’s information entropy(43) to ensure the properties contain a wealth of information that can be analyzed:

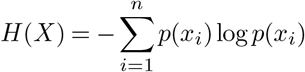

where *H*(*X*) represents the entropy of a property, *x*_*i*_ are the distinct property values, and *p*(*x*_*i*_) denotes the probability of occurrence of each value.

Based on this analysis, we selected the top three most important properties—Hydrophobicity (GRAVY), Isoelectric Point, and Surface Charge—as shown in Figure 5. These properties were standardized, with the following values for the three property:

**Fig 5.**
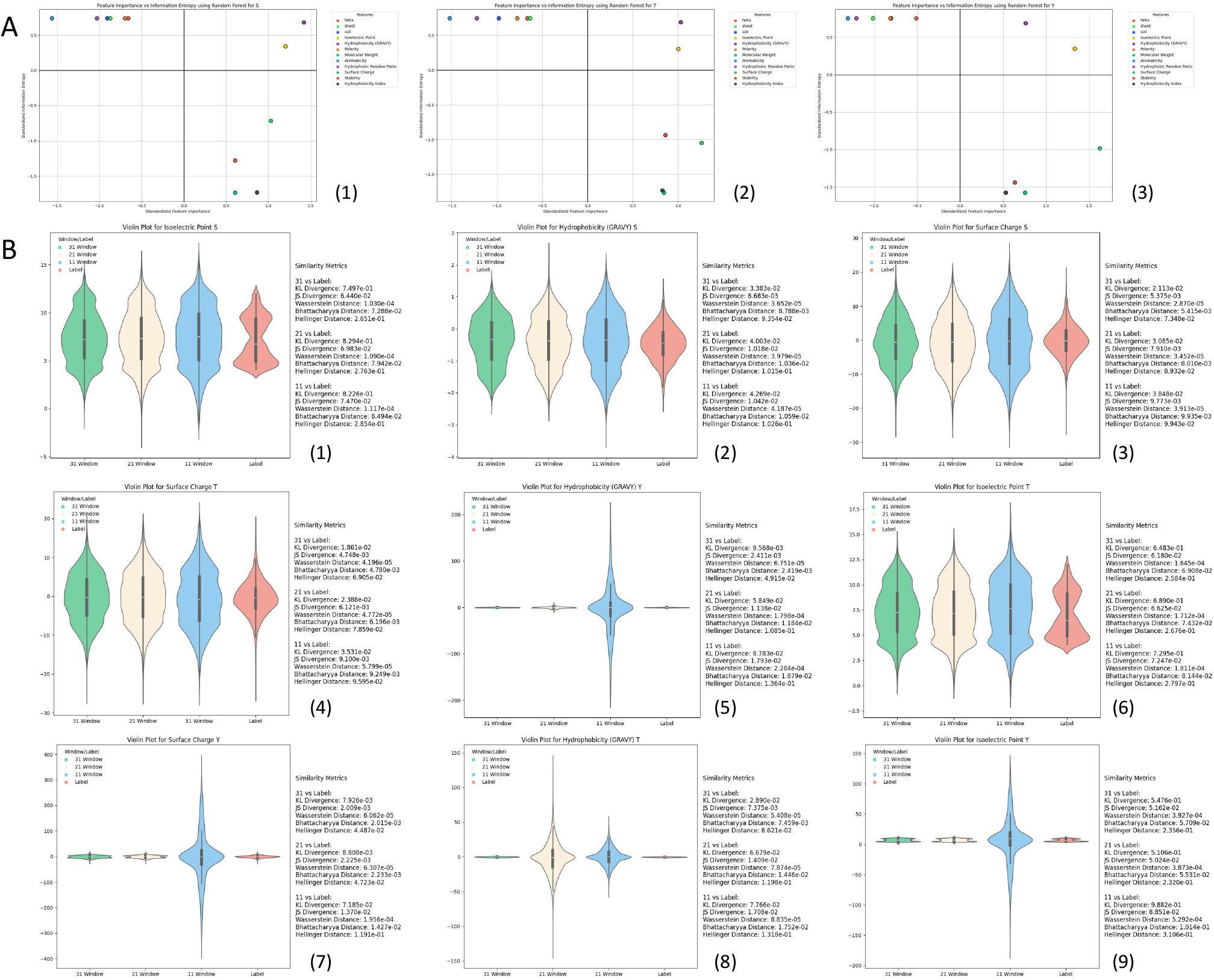
A shows the scores of important chemical properties determined by random forest for phosphorylation prediction, along with the information entropy of their distributions for Serine (S), Threonine (T), and Tyrosine (Y) sites. The data in A have been standardized, with the x-axis representing the importance of each property to phosphorylation prediction and the y-axis representing the information entropy, which indicates the amount of information contained in the property distribution. B illustrates the degree of difference between the distributions of LRT applied to window sizes of 31, 21, and 11, compared to the distributions predicted by Biopython. B also provides various metrics for comparing the distribution differences, including Kullback-Leibler (KL) divergence and Jensen-Shannon (JS) divergence. Higher similarity between distributions indicates better prediction performance and more accurate feature extraction.

- **Hydrophobicity (GRAVY)**:
  - Importance: S:0.743, T:0.752, Y:0.740
  - Entropy: S:0.376, T:0.348, Y:0.348
- **Isoelectric Point**:
  - Importance: S:0.744, T:0.752, Y:0.741
  - Entropy: S:0.380, T:0.349, Y:0.349
- **Surface Charge**:
  - Importance: S:0.744, T:0.753, Y:0.742
  - Entropy: S:0.377, T:0.348, Y:0.348

The standardization formula is:

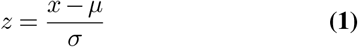

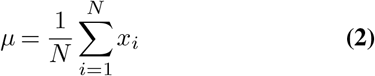

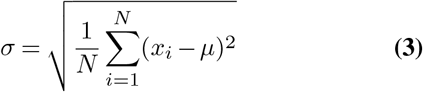

We then took 128 boundary values from the distribution of each property and paired them with the corresponding protein segments. The representations for these protein segments were extracted from the LoRA fine-tuned ESM2 model using embedding windows of sizes 31, 21, and 11, respectively. We applied multivariate linear regression to identify property directions within these embeddings.

Subsequently, all protein samples were projected onto the identified property directions, and we derived their distributions. Finally, we compared these distributions to those predicted by Biopython (Figure 5) and computed the distribution differences.

The results highlight the effectiveness of LRT in analyzing the model’s ability to extract meaningful features. By leveraging LRT, we identified property directions that are highly representative, with larger window sizes yielding more accurate and consistent distributions. These directions closely align with the predictions obtained from Biopython, demonstrating that the model’s representations successfully capture relevant physicochemical properties. This validation not only confirms the effectiveness of LRT as an analytical tool but also underscores the strong feature extraction capability of our model, showcasing its ability to capture and leverage important information for accurate predictions.

### E. Analyzing the Reasons for Suboptimal Predictive Capabilities

To investigate the reasons behind the model’s insufficient predictive capabilities, we hypothesized that similarities among phosphorylation sites might contribute to the model’s difficulties in distinguishing between them. To validate this conjecture, we conducted an experiment on the general test dataset.

We computed the representation matrices of two proteins fine-tuned with ESM2 and performed matrix subtraction. We then calculated the Frobenius norm(44) ||*A*||_*F*_ of the contrast matrix as a measure of the difference between the two proteins:

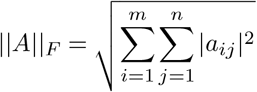

where: ||*A*|| _*F*_ is the Frobenius norm of matrix *A, m* is the number of rows in the matrix, *n* is the number of columns in the matrix, *a*_*ij*_ represents the element in the *i*-th row and *j*-th column of matrix *A*.

Since the representation matrices capture rich abstract features, not only sequence similarity but also chemical properties and structural similarities would be accounted for. We selected the two proteins with the low norms, identified by their IDs and positions, and analyzed their sequences and structures (see Figure 6). This confirms the validity of measuring differences in this manner.

**Fig 6.**
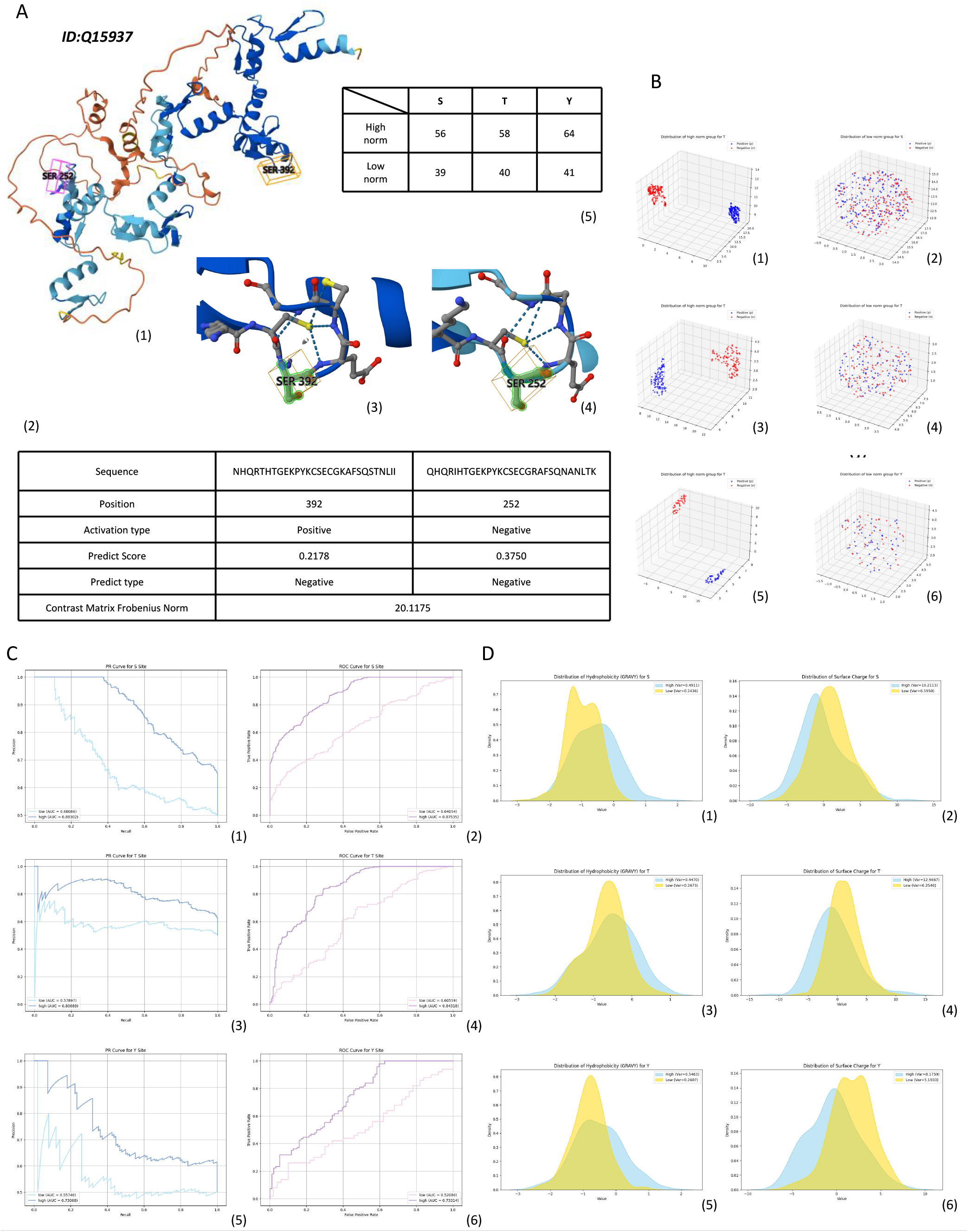
A presents a set of cases with lower and upper bounds of norm values for the high and low norm groups respectively, accompanied by a structural diagram. The norm information indicates that the protein differences within the high norm group will not be lower than the lower bound, and vice versa for the low norm group. B shows the distribution of the high and low norm groups after dimensionality reduction to 3D using UMAP. C presents the PR and ROC curves for the model’s predictions on the high and low norm groups. D shows the density distribution and variance of the properties Hydrophobicity (GRAVY) and Isoelectric Point predicted by Biopython for the high and low norm groups.

Next, we divided the 10% general test dataset into two parts based on their Frobenius norms: a high-norm group and a low-norm group. The differences in norms between any negative and positive samples in the high-norm group were set to be greater than or equal to a dynamically determined threshold (approximately allowing 0.4% of the original sample size in each group). The low-norm group was handled similarly. The specific data obtained is as follows:

- **Serine (S) Sites:**
  - High-norm group: *F >* 56 (positive: 886, negative: 171)
  - Low-norm group: *F <* 39 (positive: 192, negative: 502)
- **Threonine (T) Sites:**
  - High-norm group: *F >* 58 (positive: 365, negative: 111)
  - Low-norm group: *F <* 40 (positive: 109, negative: 500)
- **Tyrosine (Y) Sites:**
  - High-norm group: *F >* 64 (positive: 183, negative: 44)
  - Low-norm group: *F <* 41 (positive: 50, negative: 139)

Subsequently, we balanced the dataset by reducing the larger group, facilitating the subsequent experiments.

The classification results indicated a clear distinction between the two groups. To further visualize the effectiveness of this distinction, we employed 3D UMAP, which provided an intuitive representation of the data (see Figure 6).

Subsequently, we performed predictions on both norm groups. The results showed significant differences in performance, highlighting that the high-norm group had markedly better predictive capabilities compared to the low-norm group(see in 6).

Further analysis of the distribution of chemical properties revealed notable variance in the density plots(see in 6), validating our hypothesis that the similarity among certain phosphorylation sites could hinder the model’s ability to make accurate predictions. This suggests that the quality of the dataset plays a crucial role in model performance.

In conclusion, our findings indicate that the similarities among certain phosphorylation sites like chemical properties, structures significantly impact predictive capabilities.

### F. Evaluation of PhosF3C across Various Protein Tasks

To evaluate the generalization ability of the PhosF3C model across various protein-related tasks, we conducted experiments on three significant bioactivities: histone lysine crotonylation (Kcr)(45), methylation, and the Sequential and Spatial Methylation Fusion Network (SSMFN)(46). Each task represents distinct biological challenges. Kcr prediction focuses on identifying histone lysine crotonylation sites, which play key roles in cellular regulation and are implicated in various human diseases. Methylation prediction involves detecting glutamine methylation sites, a process crucial for gene regulation and cancer progression. The SSFMN task, designed to predict protein methylation sites, leverages neural networks to improve the efficiency and accuracy of post-translational modification site identification.

The model’s performance was evaluated using several important metrics. The results highlight that PhosF3C demonstrates significant improvements over other models in specific tasks. For example, in the SSMFN task, PhosF3C outperforms with an AUC of 0.9128 and accuracy of 0.8308, compared to SSMFN’s AUC of 0.9011 and accuracy of 0.8288. In the Deep-Kcr task, PhosF3C also excels, achieving an AUC of 0.8989 and accuracy of 0.8225, surpassing Deep-Kcr’s AUC of 0.8698 and accuracy of 0.7364. These results confirm that PhosF3C not only excels in individual tasks but also showcases robust generalization across a wide range of protein-related predictions. More details about the model performance were shown in Supplementary.

## 4: Conclusion

In this study, we proposed a novel framework for phosphorylation site prediction that integrates CNN and transformer architectures to capture both local and global sequence features. By using ESM2 embeddings, our model effectively represents protein sequences, enabling robust phosphorylation site prediction with state-of-the-art performance.

We also introduced FCU to enhance feature interaction and further improve model performance. Additionally, we applied LRT to evaluate the model’s feature extraction capabilities. The results showed that the model captures relevant biochemical signals, with larger window sizes providing a better representation of physicochemical distributions.

Contributions Efficient Feature Extraction and Prediction: We developed a model that uses LoRA fine-tuning and the Conformer architecture. This combination enables efficient feature extraction, accurate prediction of short protein segments, and significant reductions in both storage and computational resource requirements.

High Generalizability: Our framework has been successfully applied to various protein-related tasks, demonstrating its versatility and adaptability across different protein prediction challenges. This highlights its reliability for broader applications in bioinformatics.

Interpretability Enhancement: We introduced a top-down analysis method called LRT to examine protein representations derived from ESM2 embeddings. This approach employs traditional machine learning techniques to identify the principal directions of properties within the representations, providing valuable insights that enhance model interpretability.

However, challenges remain. One key issue is the imbalance in datasets, which can introduce biases in large-scale protein analysis. Current sampling strategies may not fully address this concern. Additionally, the reliance on high-quality sequence embeddings limits the applicability of the model to incomplete protein sequences. Future work should focus on incorporating more experimental data, improving interpretability, and extending the framework to support multimodal inputs, such as structural or interaction data.

In conclusion, our framework advances phosphorylation site prediction and holds potential for broader biological insights. Its applications in drug development and personalized medicine make it a significant step forward in computational biology and AI-driven research.

## Supporting information

This is the supplementary file of PhosF3C

