## Supplementary material for "PhosF3C: A Feature Fusion Architecture with Fine-Tuned Protein Language Model and Conformer for prediction of general phosphorylation site": This is the supplementary file of PhosF3C

**This PDF file includes:**

Supplementary Text For Experiments

Figures S1 to S4

Tables S1 to S4

### Supplementary Text For Experiments

#### Model performance on task-specific datasets

As demonstrated in supplementary Figure S1,S2 and Table S1,S2, the PhosF3C model consistently outperforms other methods on both the PhosAF and DeepIPs datasets, showcasing its robust feature extraction capabilities and suitability for phosphorylation site prediction tasks.

#### Details about the importance value and entropy of different properties

Using Random Forest and Biopython to analyze the importance of different chemical properties and the distribution of information entropy, as shown in Table S3,S4

#### Additional chemical properties' distribution on low and high norm group

We used Biopython to calculate various biochemical properties, including Isoelectric Point, Hydrophobicity (GRAVY), Polarity, Molecular Weight, Aromaticity, Hydrophobic Residue Ratio, Stability, and Hydrophobicity Index. Figure S3 shows the distribution of various chemical properties in high and low F-norm groups across S, T, and Y residues. Based on the specific data, the following characteristics can be observed:

- **Isoelectric Point**: The distribution of the high F-norm group is broader, with a standard deviation of **1.12**, while the low F-norm group has a smaller standard deviation of **0.87**, indicating greater variability in the high F-norm group.
- **Molecular Weight**: The peak of the high F-norm group is flatter, with a standard deviation of **412.6**, compared to the low F-norm group's standard deviation of **298.4**, suggesting that the high F-norm group contains more extreme values.
- **Hydrophobicity Index**: The high F-norm group exhibits a wider range with a larger tail, having a standard deviation of **4.75**, while the low F-norm group's standard deviation is only **3.25**.

For the Frobenius norm groups, the high-norm group displayed a broader and more dispersed distribution, indicating greater variability in these biochemical properties. In contrast, the low-norm group exhibited a more compact and consistent distribution. This difference highlights the presence of richer abstract feature information in the high-norm group, suggesting its potential to capture diverse physicochemical and structural characteristics.

### Other Protein Task

In this work, we present a comparison of dataset and model performance across three different protein-related tasks: DeepKCR, Methylation, and SSMFN. see in Figure S4

### Training Hyperparameters

The training process utilizes LoRA (Low-Rank Adaptation) for efficient fine-tuning of the model, reducing memory usage while allowing parameter updates. For the first configuration, a batch size of 256 is employed, and the Adam optimizer is used with a learning rate of $1 \times 10^{-4}$, betas set to $\beta_1 = 0.9$, $\beta_2 = 0.999$, and a weight decay of $1 \times 10^{-4}$. The second configuration, which uses the Conformer architecture, applies a smaller batch size of 64, with the Adam optimizer configured with a learning rate of $5 \times 10^{-5}$, betas of $\beta_1 = 0.9$, $\beta_2 = 0.999$, and a weight decay of $1 \times 10^{-4}$. The loss function for both configurations is Cross-Entropy Loss. The model architecture includes both a Transformer branch and a CNN branch, where the weights for both branches are set equally at 0.5, ensuring balanced contributions from each branch in the final output and early stopping were applied to handle the problem of overfitting.

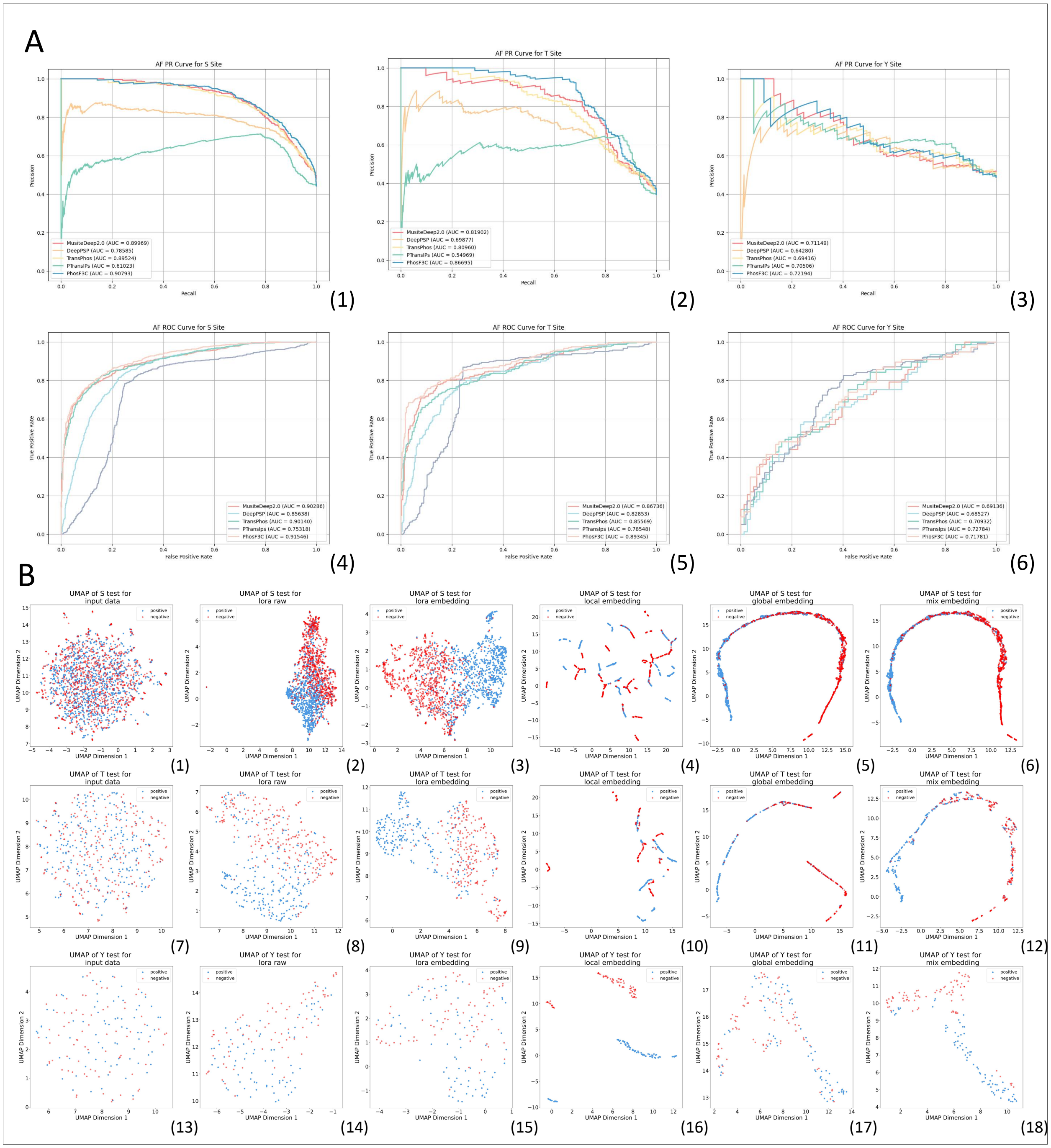


**Figure S1**: A presents the PR and ROC curves for performance evaluation on PhosAF Dataset, B shows the UMAP visualization of the dataset, providing a low-dimensional representation of the data distribution during training.

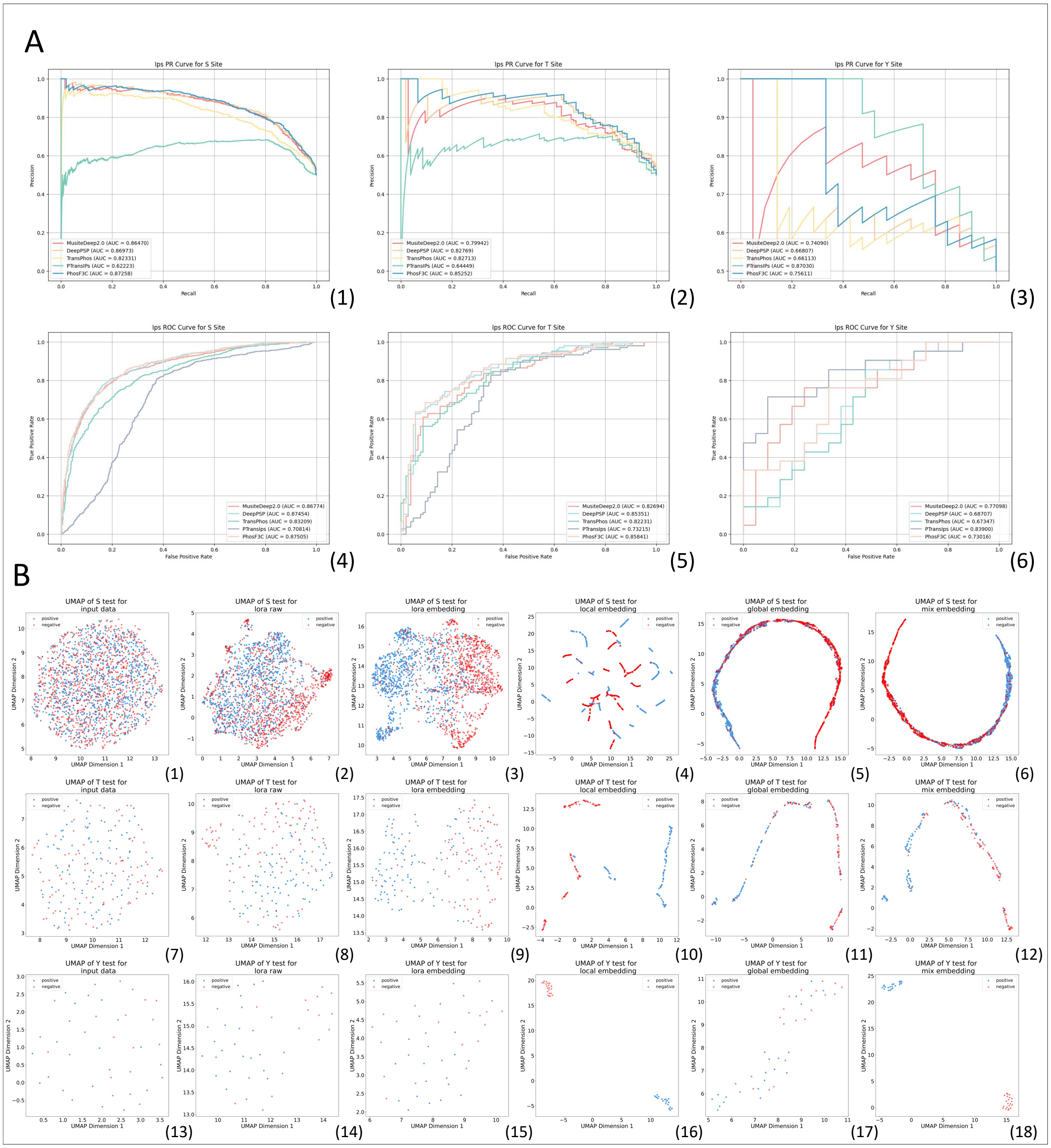


**Figure S2**: A presents the PR and ROC curves for performance evaluation on DeepIps Dataset, B shows the UMAP visualization of the dataset, providing a low-dimensional representation of the data distribution during training.

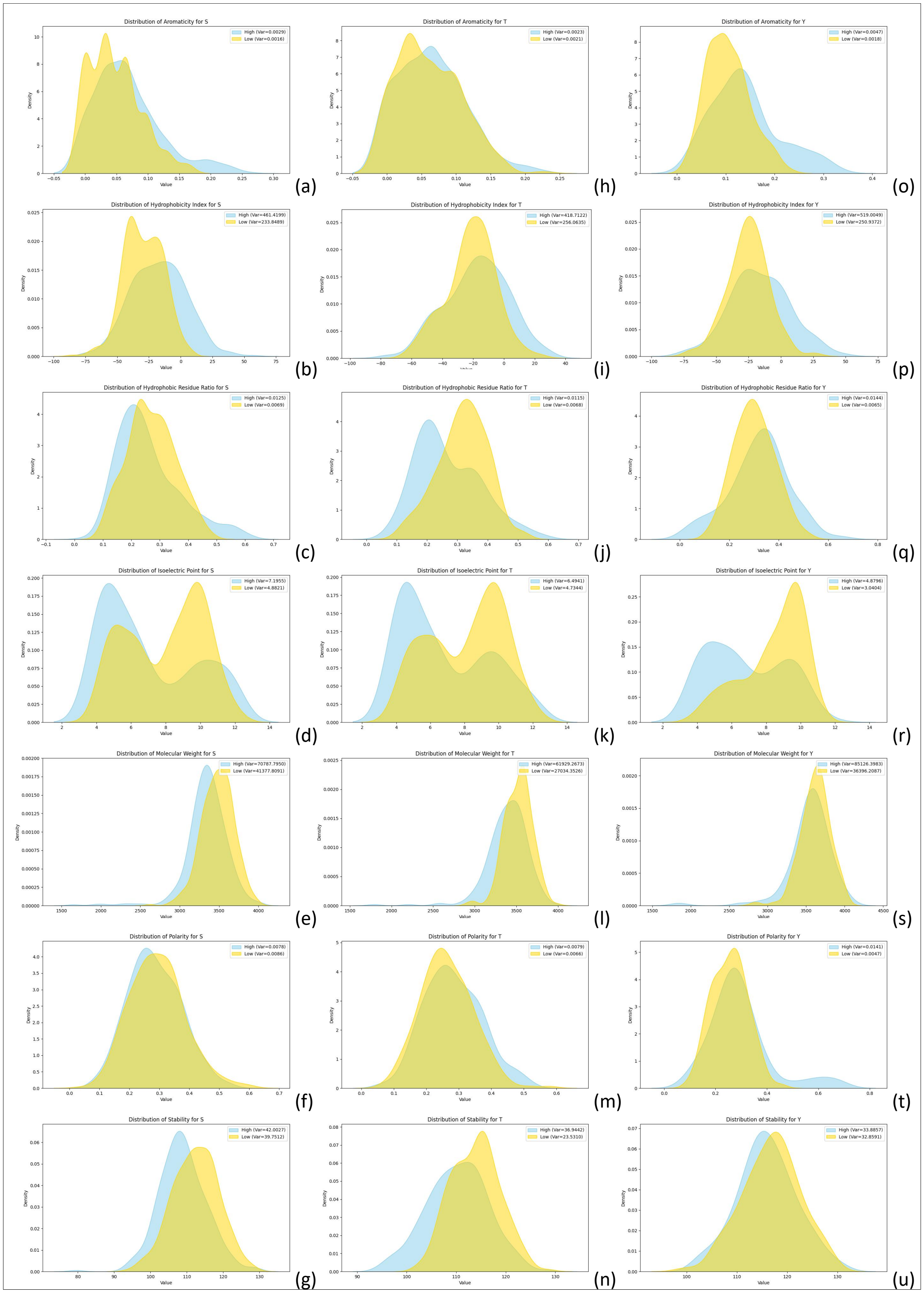


**Figure S3**: Distribution density of all properties in the high and low norm groups, with the variances of the distributions also recorded to capture the spread and variability within each group.

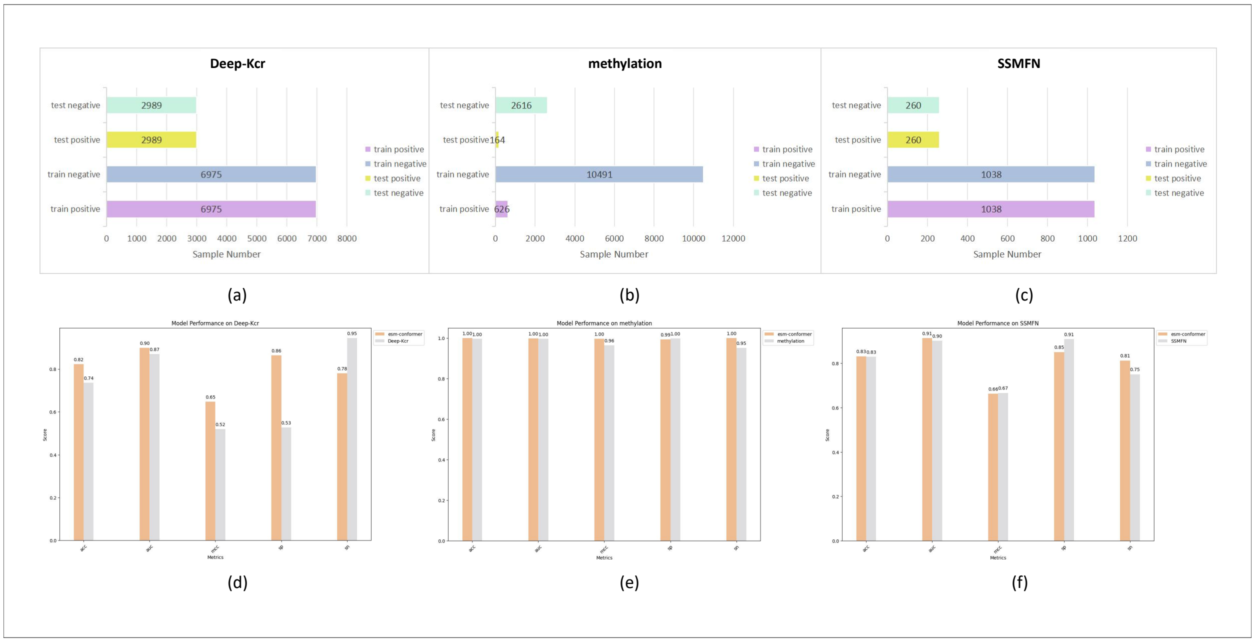


**Figure S4**: Figures (a), (b), and (c) display the data information for three protein-related tasks: histone lysine crotonylation (Kcr), methylation, and the Sequential and Spatial Methylation Fusion Network (SSMFN), along with the distribution of positive and negative samples for each dataset. Figures (d), (e), and (f) show the comparison of the PhosF3C model's performance with baseline models in these tasks.

| Type | Model | AUC | ACC | MCC | F1 | RECALL | PRECISION |
|---|---|---|---|---|---|---|---|
| S | PSP | 0.8564 | 0.7731 | 0.5628 | 0.7702 | 0.8601 | 0.6974 |
| | Musite | 0.9029 | 0.789 | 0.5965 | 0.7874 | 0.8834 | 0.7102 |
| | Phos | 0.9014 | 0.7869 | 0.5914 | 0.7847 | 0.8785 | 0.709 |
| | IPS | 0.7532 | 0.7555 | 0.5245 | 0.7503 | 0.8309 | 0.684 |
| | PhosF3C | 0.9155 | 0.8264 | 0.6542 | 0.813 | 0.8533 | 0.7763 |
| T | PSP | 0.8285 | 0.7719 | 0.526 | 0.6996 | 0.7796 | 0.6346 |
| | Musite | 0.8674 | 0.7928 | 0.5696 | 0.7256 | 0.8041 | 0.6611 |
| | Phos | 0.8557 | 0.7872 | 0.5452 | 0.7086 | 0.7592 | 0.6643 |
| | IPS | 0.7855 | 0.7608 | 0.5516 | 0.7162 | 0.8857 | 0.6011 |
| | PhosF3C | 0.8934 | 0.7914 | 0.5733 | 0.7292 | 0.8245 | 0.6537 |
| Y | PSP | 0.6853 | 0.6139 | 0.2311 | 0.6258 | 0.6623 | 0.593 |
| | Musite | 0.6914 | 0.6139 | 0.239 | 0.6474 | 0.7272 | 0.5833 |
| | Phos | 0.7093 | 0.6456 | 0.3077 | 0.6818 | 0.7792 | 0.606 |
| | IPS | 0.7278 | 0.69 | 0.399 | 0.7232 | 0.8265 | 0.6429 |
| | PhosF3C | 0.7178 | 0.6329 | 0.2872 | 0.6778 | 0.7922 | 0.5922 |

**Table S1**: Performance metrics on the PhosAF dataset categorized by phosphorylation site type.

| Type | Model | AUC | ACC | MCC | F1 | RECALL | PRECISION |
|---|---|---|---|---|---|---|---|
| S | PSP | 0.8745 | 0.7464 | 0.5169 | 0.7797 | 0.8973 | 0.6893 |
| | MusiteDeep2.0 | 0.8677 | 0.731 | 0.4972 | 0.773 | 0.9158 | 0.6687 |
| | Phos | 0.8321 | 0.7012 | 0.4384 | 0.7506 | 0.8994 | 0.6441 |
| | IPS | 0.7081 | 0.7928 | 0.425 | 0.7414 | 0.8522 | 0.6561 |
| | PhosF3C | 0.875 | 0.751 | 0.5279 | 0.7843 | 0.9055 | 0.6918 |
| T | PSP | 0.8535 | 0.7714 | 0.5453 | 0.7818 | 0.819 | 0.7478 |
| | MusiteDeep2.0 | 0.8269 | 0.7524 | 0.5123 | 0.7719 | 0.8381 | 0.7154 |
| | Phos | 0.8223 | 0.7333 | 0.4709 | 0.75 | 0.8 | 0.7059 |
| | IPS | 0.7322 | 0.7095 | 0.4386 | 0.7469 | 0.8571 | 0.6618 |
| | PhosF3C | 0.8584 | 0.7667 | 0.5466 | 0.7897 | 0.8762 | 0.7188 |
| Y | PSP | 0.6871 | 0.6429 | 0.286 | 0.6512 | 0.6667 | 0.6364 |
| | MusiteDeep2.0 | 0.771 | 0.7381 | 0.4767 | 0.7442 | 0.7619 | 0.7273 |
| | Phos | 0.6735 | 0.619 | 0.2392 | 0.6364 | 0.6667 | 0.6087 |
| | IPS | 0.839 | 0.7143 | 0.4472 | 0.75 | 0.8571 | 0.6667 |
| | PhosF3C | 0.7302 | 0.6429 | 0.2942 | 0.6809 | 0.7619 | 0.6154 |

**Table S2**: Performance metrics on the DeepIps dataset categorized by phosphorylation site type.

| Feature | S | T | Y |
|---|---|---|---|
| **Helix** | -0.977 | -0.991 | -0.994 |
| **Sheet** | -1.098 | -1.089 | -1.210 |
| **Coil** | -1.086 | -0.997 | -1.193 |
| **Isoelectric Point** | 0.731 | 0.904 | 0.952 |
| **Hydrophobicity (GRAVY)** | 1.362 | 1.452 | 0.952 |
| **Polarity** | -0.628 | -0.840 | -0.457 |
| **Molecular Weight** | 0.537 | 0.780 | 1.212 |
| **Aromaticity** | -1.170 | -1.312 | -1.155 |
| **Hydrophobic Residue Ratio** | -0.747 | -0.629 | -0.813 |
| **Surface Charge** | 1.010 | 0.994 | 1.227 |
| **Stability** | 0.530 | 0.864 | 0.607 |
| **Hydrophobicity Index** | 1.536 | 0.864 | 0.875 |

**Table S3**: Feature inportance values for Serine (S), Threonine (T), and Tyrosine (Y) across various properties.

| Feature | S | T | Y |
|---|---|---|---|
| **Helix** | 0.744 | 0.752 | 0.741 |
| **Sheet** | 0.743 | 0.752 | 0.741 |
| **Coil** | 0.744 | 0.752 | 0.741 |
| **Isoelectric Point** | 0.381 | 0.349 | 0.349 |
| **Hydrophobicity (GRAVY)** | 0.705 | 0.716 | 0.697 |
| **Polarity** | 0.744 | 0.753 | 0.742 |
| **Molecular Weight** | -1.680 | -1.737 | -1.760 |
| **Aromaticity** | 0.745 | 0.753 | 0.742 |
| **Hydrophobic Residue Ratio** | 0.744 | 0.752 | 0.741 |
| **Surface Charge** | -1.107 | -1.181 | -1.189 |
| **Stability** | -1.087 | -1.237 | -0.858 |
| **Hydrophobicity Index** | -1.675 | -1.424 | -1.688 |

**Table S4**: Feature entropy values for Serine (S), Threonine (T), and Tyrosine (Y) across various properties.
